# Optimising passive surveillance of a neglected tropical disease in the era of elimination: A modelling study

**DOI:** 10.1101/2020.07.20.211714

**Authors:** Joshua Longbottom, Charles Wamboga, Paul R. Bessell, Steve J. Torr, Michelle C. Stanton

**Affiliations:** Department of Vector Biology, Liverpool School of Tropical Medicine, Liverpool, L3 5QA; Centre for Health Informatics, Computing and Statistics, Lancaster Medical School, Lancaster University, Lancaster, LA1 4YW; Ministry of Health, Kampala, Uganda; Epi Interventions, Edinburgh, UK

## Abstract

**Background:** Surveillance is an essential component of global programs to eliminate infectious diseases and avert epidemics of (re-)emerging diseases. As the numbers of cases decline, costs of treatment and control diminish but those for surveillance remain high even after the ‘last’ case. Reducing surveillance may risk missing persistent or (re-)emerging foci of disease. Here, we use a simulation-based approach to determine the minimal number of passive surveillance sites required to ensure maximum coverage of a population at-risk (PAR) of an infectious disease.

**Methodology and Principal Findings:** For this study, we use Gambian human African trypanosomiasis (g-HAT) in north-western Uganda, a neglected tropical disease (NTD) which has been reduced to historically low levels (<1000 cases/year globally), as an example. To quantify travel time to diagnostic facilities, a proxy for surveillance coverage, we produced a high spatial-resolution resistance surface and performed cost-distance analyses. We simulated travel time for the PAR with different numbers (1-170) and locations (170,000 total placement combinations) of diagnostic facilities, quantifying the percentage of the PAR within 1h and 5h travel of the facilities, as per in-country targets. Our simulations indicate that a 70% reduction (51/170) in diagnostic centres still exceeded minimal targets of coverage even for remote populations, with >95% of a total PAR of ~3million individuals living ≤1h from a diagnostic centre, and we demonstrate an approach to best place these facilities, informing a minimal impact scale back.

**Conclusions:** Our results highlight that surveillance of g-HAT in north-western Uganda can be scaled back without substantially reducing coverage of the PAR. The methodology described can contribute to cost-effective and equable strategies for the surveillance of NTDs and other infectious diseases approaching elimination or (re-)emergence.

**Author Summary:** Disease surveillance systems are an essential component of public health practice and are often considered the first line in averting epidemics for (re-)emerging diseases. Regular evaluation of surveillance systems ensures that they remain operating at maximum efficiency; systems that survey diseases of low incidence, such as those within elimination settings, should be simplified to reduce the reporting burden. A lack of guidance on how to optimise disease surveillance in an elimination setting may result in added expense, and/or the underreporting of disease. Here, we propose a framework methodology to determine systematically the optimal number and placement of surveillance sites for the surveillance of infectious diseases approaching elimination. By utilising estimates of geographic accessibility, through the construction of a resistance surface and a simulation approach, we identify that the number of operational diagnostic facilities for Gambian human African trypanosomiasis in north-western Uganda can be reduced by 70% with a minimal reduction in existing coverage, and identify the minimum number of facilities required to meet coverage targets. Our analysis can be used to inform the number and positioning of surveillance sites for diseases within an elimination setting. Passive surveillance becomes increasingly important as cases decline and active surveillance becomes less cost-effective; methods to evaluate how best to engage this passive surveillance capacity given facility capacity and geographic distribution are pertinent for several NTDs where diagnosis is complex. Not only is this a complicated research area for diseases approaching elimination, a well-designed surveillance system is essential for the detection of emerging diseases, with this work being topical in a climate where emerging pathogens are becoming more commonplace.

## Introduction

Quantifying the spatial and temporal distribution of cases is essential for the development, implementation and monitoring of programmes to control or eliminate infectious diseases (1–3). In general, costs for treatment and control of a disease scale with the number of cases, but those for surveillance can be high even when disease incidence is low (4). Programmes aiming to eliminate or eradicate a disease maintain high levels of surveillance, as illustrated by the global programmes against smallpox and polio (5, 6). Similarly, surveillance systems to detect (re-)emerging diseases may also have relatively large surveillance costs in relation to the numbers of cases (7). Maintaining adequate surveillance for diseases approaching elimination or with potential for emergence can be prohibitively costly for national health systems, particularly for resource-poor countries in the tropics (8, 9), and as such, approaches for cost-effective surveillance are required.

Neglected tropical diseases (NTDs) cause high levels of mortality and morbidity in some of the world’s poorest countries (10, 11). A global program aims to eliminate or eradicate eleven NTDs by 2030 and as the elimination of several NTDs approaches, globally and/or nationally, there is a pressing need for cost-effective strategies to detect the last remaining cases (12, 13). The new surveillance strategies will need to be adapted to the disease itself, local health systems and the population at risk (PAR) (1).

Disease surveillance systems include active, passive, sentinel, and statistical approaches, but the majority of reportable disease surveillance is conducted through passive surveillance (14, 15). Passive surveillance is based primarily on compilation of case reports submitted by healthcare facilities, produced when infected individuals present to the facility for diagnosis and treatment. One approach to improving cost-effectiveness of passive surveillance is to quantify the minimal number of surveillance sites that provides maximal coverage of the PAR. Towards this aim, we combined estimates of travel time to passive surveillance sites with a simulation-based approach to determine the optimal number of sites required to ensure that 50% and 95% of a PAR of Gambian human African trypanosomiasis (g-HAT) are within 1-hour and 5-hours of a diagnostic facility, respectively. The aforementioned criteria come from in-country targets (16), however, the approach described here can be adapted for a range of NTDs in an elimination setting, with varying coverage and travel time requirements determined by national governments, NGOs or other disease surveillance operators.

### Scale back in the context of human African trypanosomiasis

Human African trypanosomiasis (HAT, also called ‘sleeping sickness’) is an NTD occurring across sub-Saharan Africa, caused by sub-species of *Trypanosoma brucei* transmitted by tsetse flies. Most (>95%) cases of HAT are caused by *T. b. gambiense* (g-HAT), for which humans are the primary host; the remaining ~5% of cases are caused by *T. b. rhodesiense* (r-HAT) for which the primary reservoirs are animal hosts (17). No prophylaxis or vaccine exists for either form of HAT. Therapeutic drugs are available (18), but the toxic nature of chemotherapy requires that infection status is confirmed before the patient is treated. Disease prevention and control efforts focus on the detection and treatment of existing cases and control of the tsetse vector (19).

The World Health Organization (WHO) aims to eliminate g-HAT as a public health problem by 2020. The definitions of elimination include fewer than 2,000 cases reported per annum globally, a 90% reduction in areas reporting >1 case in 10,000 per annum compared to 2000-2004, with a country-level indicator of elimination defined as fewer than 1 reported case per 10,000 people, per annum (averaged over a 5-year period) in each health district, in conjunction with adequate, functional control and surveillance (20). Uganda is on track to achieve this goal with a reduction from 2,757 (range, 310-948 cases/year) cases reported between 2000-2004 to 18 (range, 0-9 cases/year) cases in 20014-18 (21). These reductions are mostly attributable to active screening and treatment of the population (22, 23), and, more recently, vector control (24). For g-HAT, active screening has been the cornerstone of surveillance in most foci during periods of high case numbers. As case numbers decline this has become increasingly less cost-effective and surveillance has switched to a passive system.

Plans to scale back the number of passive surveillance sites operating RDTs are ongoing in Uganda, with the target of ensuring that by 2030, ≥50% of at-risk populations should live within 1-hour of a health facility with HAT diagnostics and ≥95% should live within 5-hours (16). The number of health facilities using HAT RDTs were reviewed in July 2014 and September 2015, in response to the evolving epidemiological situation of g-HAT in Uganda (25). Previous analyses utilised the Euclidean distance to facilities as opposed to estimates of travel time. The number of facilities in operation may be further reviewed to better quantify accessibility, identify the minimal number required to ensure sufficient coverage, and to maximise cost-effectiveness.

Optimisation of passive screening capacity in NW Uganda must consider two key elements. First, there must be evidence of reduced transmission, be this supported by active screening or reduced abundance of tsetse (23, 24). Second, g-HAT has a very long interval between infection and detection of a case, and cases can remain undetected for some time (26), therefore we are required to consider the long tail of this case distribution and any reservoirs of infection (27). Consequently, the resource review proposed here should target a wider area that has been historically at risk in order to detect any residual infections in the area. The streamlining or optimisation of the number of facilities might be considered a step towards establishment of a network of sentinel screening sites to monitor post-elimination (17).

The work described here aims to utilise information on the location of operational HAT diagnostic facilities, alongside estimates of travel time to said facilities, to inform an analysis designed to identify the optimal number, and placement of, surveillance facilities required to meet the in-country coverage targets defined above.

## Methods

### Study setting

The focal area of this study is north-western Uganda where we focus on seven districts within the West Nile region which form Uganda’s g-HAT endemic area. These endemic districts are Arua (2,100 km^2^), Maracha (693 km^2^), Koboko (862 km^2^), Amuru (3,625km^2^), Adjumani (3,030km^2^), Moyo (1,800km^2^), and Yumbe (1,524 km^2^). Collectively, these districts have a population of approximately 3.5 million people (28). For the purpose of this study, from this point onwards, we refer to these seven districts as “north-western Uganda”.

### High-resolution resistance surface

To quantify travel time to diagnostic facilities, we first sought to generate a resistance surface for north-western Uganda at a 30m × 30m spatial resolution. Resistance surfaces (also termed ‘friction surfaces’) contain estimates of associated travel cost for gridded cells within a Cartesian plane and are used within cost-distance analyses to quantify the effort required to travel between each cell and an origin point (29). To construct the resistance surface, we collated data from a variety of sources. We obtained comprehensive road network data from OpenStreetMap (30), and assigned travel speeds along differing road classes based off observed speeds from GPS tracks obtained during February-April 2018 (31). Speeds were representative of motorcycle travel, in agreement with the most commonly used mode of transportation in Uganda (32). Estimates of off-road travel were generated utilising remotely sensed Landsat-8 data (33), and a normalised difference vegetation index (NDVI) (34, 35), paired with information from studies detailing travel time through varying vegetation densities (36, 37). Finally, we combined off-road and on-road travel estimates to produce a comprehensive resistance surface detailing the time taken to traverse through each 30m × 30m gridded cell. We evaluated the accuracy of the resistance surface by comparing predicted travel time along 1,000 routes with times derived from Google Maps within a linear regression (38). An in-depth description of the process is provided within Supplementary File 1.

### Health facility data

In 2013, a new passive surveillance strategy was implemented in north-western Uganda, which saw the number of facilities participating in passive surveillance increase from four to 212, which was subsequently adjusted to 170 in 2017 (25). Each facility utilises either RDTs alone (39), or RDTs and microscopy techniques (40), and is supplemented by mini-anion exchange centrifugation technique (mAECT) (41), with three of these second group facilities also equipped with loop-mediated isothermal amplification (LAMP) for identifying suspected g-HAT cases (Fig. 1:A) (42). For facilities with available data, the number of RDTs used during 2017 is shown within Fig. 1:B. Facilities operating LAMP and LED diagnostics serve as referral sites for RDT positive individuals, and are representative of larger facilities with higher quality services (25).

**Figure 1.**
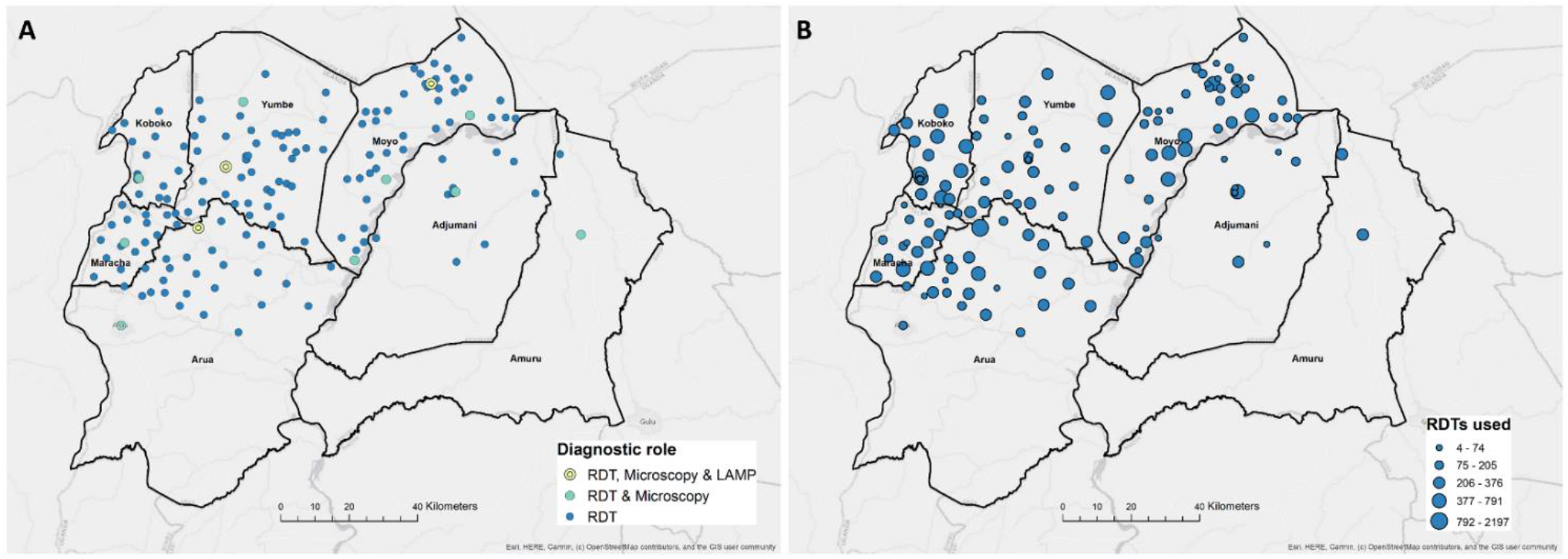
A: Location of existing HAT diagnostic centres across endemic districts in north-western Uganda (2017, *n* = 170), coloured by type of diagnostic method used. B: Number of HAT rapid diagnostic tests (RDTs) used by operational facilities during 2017.

### Human population density data

To detail the number of individuals within HAT risk areas who live within 1-hour and 5-hours travel time of a HAT diagnostic facility, we utilised a high-resolution population density surface (28). This surface matched the spatial resolution of the resistance surface generated above (30m × 30m) and was re-sampled to ensure gridded cells aligned. A crude estimate of the PAR of g-HAT within the study region was determined by creating a 10km radius buffer around reported cases between 2000 and 2018 (Fig. 2:A) (21, 25), representative of moderate risk (Fig. 2:B). Case data were obtained from the WHO HAT atlas (21), and the occurrence locations represent either the residence of the individual (data collected through active screening), or the location in which a case was diagnosed (if diagnosed during passive screening). Unfortunately, there is uncertainty surrounding the location of infection with both collection methods, and one of the advantages of buffering case data is that this approach accounts for some of the uncertainty surrounding the true location of infection. Several papers quantifying g-HAT risk similarly use a buffer around case data, albeit a more conservative distance of 30km (see Franco et al. 2014 (19), 2017 (17), 2018 (43) and 2020 (20)). The population-at-risk surfaces within Franco et al. are generated for the whole continent and we believe that, in the absence of a HAT-risk model, this approach is a good quantification of those at risk within north-western Uganda. Buffering case data identified 3,025,801 individuals living within at-risk areas (PAR provided as Supplementary File 2). The PAR surface used in this analysis captures the historic distribution of g-HAT within north-western Uganda and is therefore greater than PAR surfaces generated using only contemporary data (e.g. Simarro et al. (44)). The inclusion of historic data (2000–2018) provides higher confidence in the detection of disease and residual cases from across the area where transmission has occurred previously.

**Figure 2.**
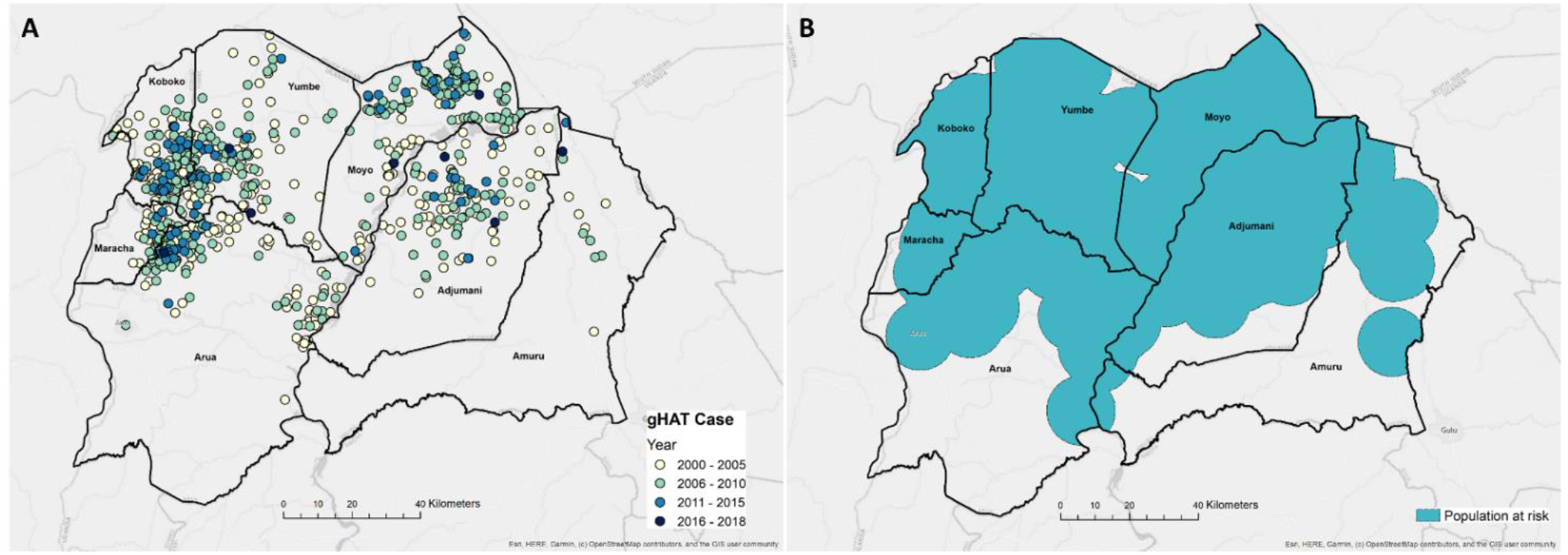
A: Spatial distribution of g-HAT cases within north-western Uganda, 2000-2018. B: Areas at risk of g-HAT, identified as locations within 10km of a case.

### Simulation of accessibility to diagnostic centres under differing placements

Information on the location of existing g-HAT diagnostic facilities may be paired with the resistance surface to answer any of the following questions:

1. What is the smallest collection of the existing facilities that would meet the coverage requirements (i.e. ≥50% of PAR within 1h; ≥95% within 5h)?
2. What reductions in health facility numbers can be made without significantly reducing current levels of coverage with respect to 1h and 5h travel time targets?
3. Given a fixed number of facilities (for example, informed by funding availability or from answering questions one or two), what is the best spatial placement of said facilities to ensure maximum coverage of the PAR?

#### Deriving the optimal number of facilities

To derive the optimal number of facilities required within north-western Uganda to ensure that ≥50% of the PAR live less than 1 hour, and ≥95% of the PAR live less than 5 hours from a health facility with HAT diagnostics respectively, we performed a simulation study. Suppose *S_n_* is the set of all possible combinations of *n* facilities from the 170 currently available. We wish to obtain an estimate of the average PAR living within 1 or 5 hours of a facility for all possible values of *n*, but once *n* > 1, the total number of combinations of *n* facilities is very large and computationally prohibitive. As such, we adopted a simulation approach, such that for *n* = 2,..,170 we generated *Ŝ_n_* which consisted of 1,000 random samples of *n* facilities. For each sample, *i* = 1,…,1000 we calculated the size of the PAR within 1, 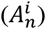 or 5, 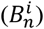 hours of any of the *n* facilities using a cost-distance analysis implemented within R (version 3.5.1). From this we can obtain *M_n_*, the average number of people within 1 hour, and *V_n_*, the average number of people within 5 hours of any of the *n* facilities (eqns. 1&2). *Q_n_* represents the proportion of simulations for which more than 50% of the PAR, (*D*), are within 1 hours travel to any of the *n* facilities (eqn. 3), and *Z_n_* represents the proportion of simulations for which more than 95% of the PAR are within 5 hours travel to any of the *n* facilities (eqn. 4). To avoid spatial clustering within samples, we utilised an inhibitory sampling approach when generating *Ŝ_n_*, ensuring a minimum distance of 4km between sampled facilities (45). The optimal facility number simulation is outlined in eqns. 1-4:

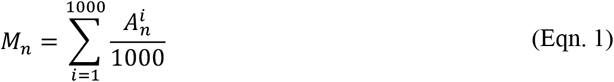

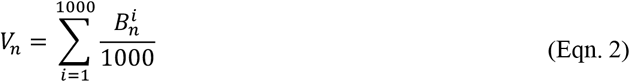

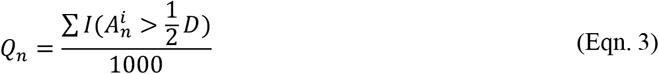

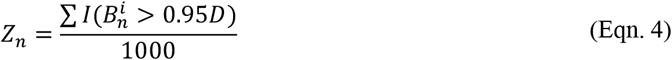

The minimal number of facilities, *N^min^*, was identified as the lowest value of *n* for which both *Q_n_* and *Z_n_* > 0.

#### Deriving the optimal placement of facilities when the facility number is known

To demonstrate how our approach can be used to determine the optimal placement of a fixed number of facilities when the target number to retain is known (i.e. informed by the above simulation, or when predefined by disease surveillance budgets), we implemented a further simulation analysis. For this simulation, when given a fixed number of facilities, *n*, we wish to identify the optimal subset of *n* facilities, from the 170 available which results in the highest percentage of the PAR within 1-hour and 5-hour travel. N’dungu et al. (46), detail in their work reporting on the Trypa-NO! project within north-western Uganda, that there is funding available to retain a maximum of 51 g-HAT diagnostic facilities from the 170 facilities currently operational. Here, using 51 facilities as an example, we demonstrate how the optimal placement of these facilities can be determined from an existing distribution using a cost-distance simulation. It should be noted that the number of available combinations (*C*(170,51)) is computationally prohibitive, therefore we employed a simulation approach utilising 10,000 random combinations of 51 facilities. For each sample, the total population within 1-hour and 5-hour of a facility was determined utilising a cost-distance analysis (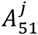 and 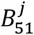 respectively, *j* = 1,…, 10000). For these simulations, we did not employ an inhibitory sampling approach, with no minimal distance enforced between selected facilities within each sample. The optimal selection of the 51 facilities was the combination that resulted in the largest number of the PAR being within the pre-defined criteria, as determined by a cumulative rank assessing percentage coverage within both 1-hour and 5-hour thresholds.

As a baseline for which to assess the impact of a scale back on accessibility of the PAR, we generated a cost-distance surface including all 170 facilities, representative of the situation in 2017. We then compared the predicted travel time for each 30 × 30m cell under the optimal scale back scenario, with the predicted travel time under the scenario representative of 2017, to determine which geographic locations would be most affected.

## Results

### High-resolution resistance surface

The high-resolution resistance surface generated showed good agreement when compared with freely obtained estimates of travel time from Google Maps, for 1,000 randomly generated validation routes across north-western Uganda (*R*^2^ = 0.902, *p* < 2e^-16^) (Fig. 3:A). The spatial distribution of the origin and destination locations used within the validation process is shown as Fig. 3:B. Whilst there is strong correlation, modelled travel times are predicted to be slightly slower when compared to Google (root-mean-square error = 14.98), this is likely due to our estimates relating to motorbike based travel, *vs* Google times being representative of car travel. Nonetheless, this validation process provided confidence in the resistance surface, and subsequently in our estimates of travel time to diagnostic facilities. To facilitate reproducibility and additional research applications within the region, the resulting resistance surface is provided as Supplementary File: 3.

**Figure 3.**
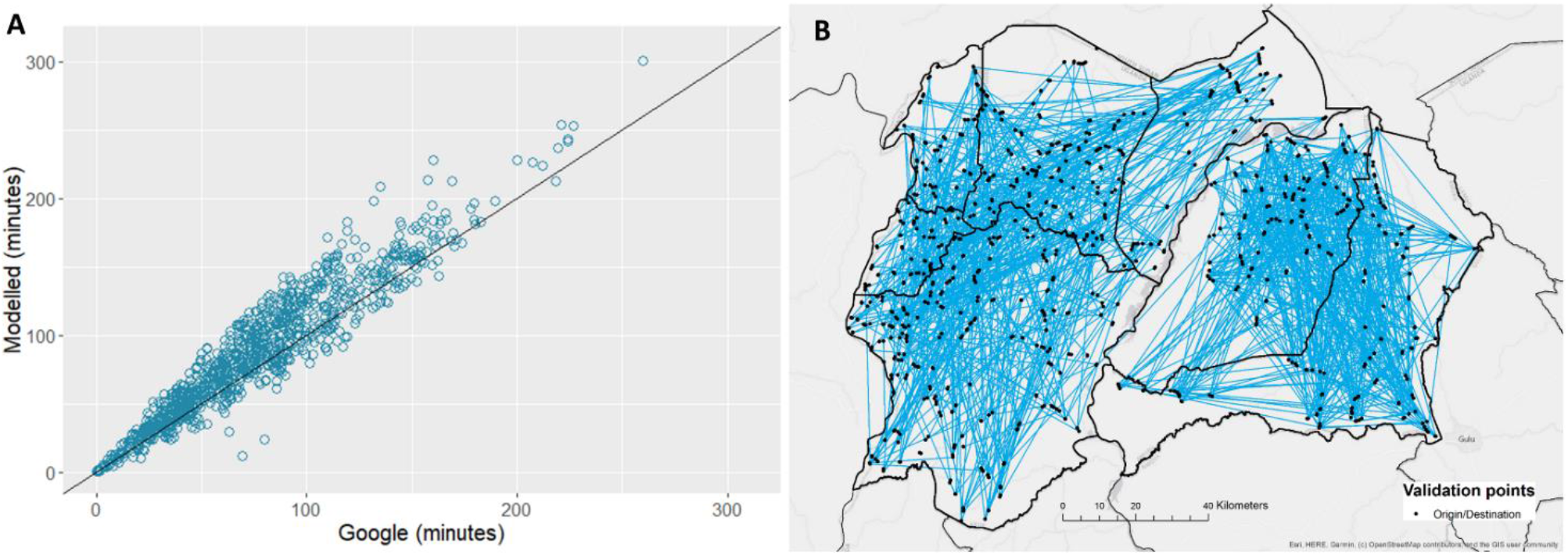
A: Result of a linear regression comparing observed travel times from Google Maps, and modelled travel time (our surface) (*R*^2^ = 0.902, *p* < 2e^−16^). B: Spatial distribution of origin/destination locations (black dots) used to validate the resistance surface. Blue lines represent pairing of origin/destination locations used to inform the regression presented in A.

### Deriving the minimal number of facilities

Results from the simulation experiment defined in Eq. 1-4 indicate that within north-western Uganda, a minimum of three facilities are required to ensure that 50% of the PAR live within 1-hours travel of a HAT diagnostic facility, and a minimum of one facility is required to ensure and 95% of the PAR live within 5-hours travel (Fig. 4), when considering a subset (1000 simulations) of all possible combinations of *n* facilities. These results, however, may be biased toward highly populated areas (i.e. urban centres) and it is unlikely that an initial scale back of this magnitude would be adopted by a national program due to a range of political and contextual factors. We present a plot showing the median percentage of the population living outside of 1-hours travel time as Fig. 4B. This plot demonstrates that to maintain similar levels of coverage to that achieved by all 170 facilities in operation, scaling back to 51 facilities would ensure that 95.25% of the PAR would be within 1-hours travel of a facility, opposed to ~50% of the PAR with the much reduced scale back to three facilities. We focus on this 1-hour threshold, as the 5-hour threshold is easily attained across our study area. Although we demonstrate that the ≥50% within 1-hour and ≥95% within 5-hour coverage targets can be achieved with very few facilities, our results question the effects of such a dramatic scale back on access inequality. Coverage targets can be easily met by a low number of facilities if such facilities are located within urban centres; this approach, however, may further isolate remote rural populations if care isn’t taken when planning a scale back.

**Figure 4.**
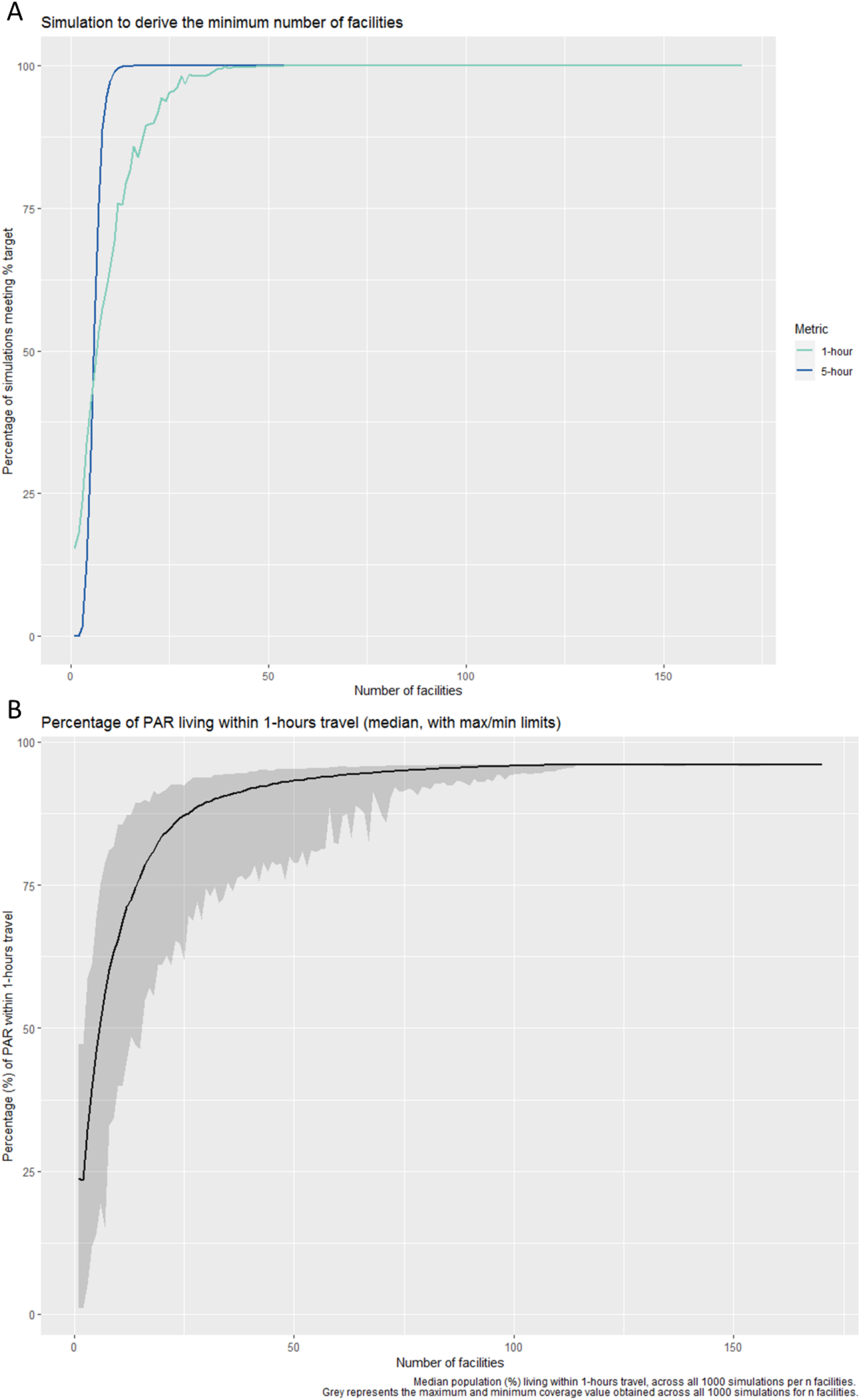
A: Results from the simulation to derive the minimal number of facilities required to ensure 50% of the population live within 1-hour travel time of a facility with HAT diagnostics (green) and 95% of the population live within 5-hours travel time of a facility with HAT diagnostics (blue). Results are shown as the percentage of simulations (from a total of 1000) which meet the ≥50% and ≥95% coverage targets. B: Median population (expressed as a percentage) living outside of 1-hours travel, across all 1000 simulations per *n* facilities. Grey represents the maximum and minimum % coverage value obtained across all 1000 simulations per *n* facilities.

### Deriving the optimal placement of facilities when the facility number is known

As a baseline for which to assess the impact of a scale back on the accessibility of the PAR, we first generated estimates of accessibility to all 170 facilities which were operational in 2017 (Fig. 5:A-B). The results from the cost-distance analysis utilising all 170 facilities indicate that 96.15% and 99.77% of the population at risk of HAT live within 1-hour and 5-hours travel time of a HAT diagnostic facility in 2017 respectively.

**Figure 5.**
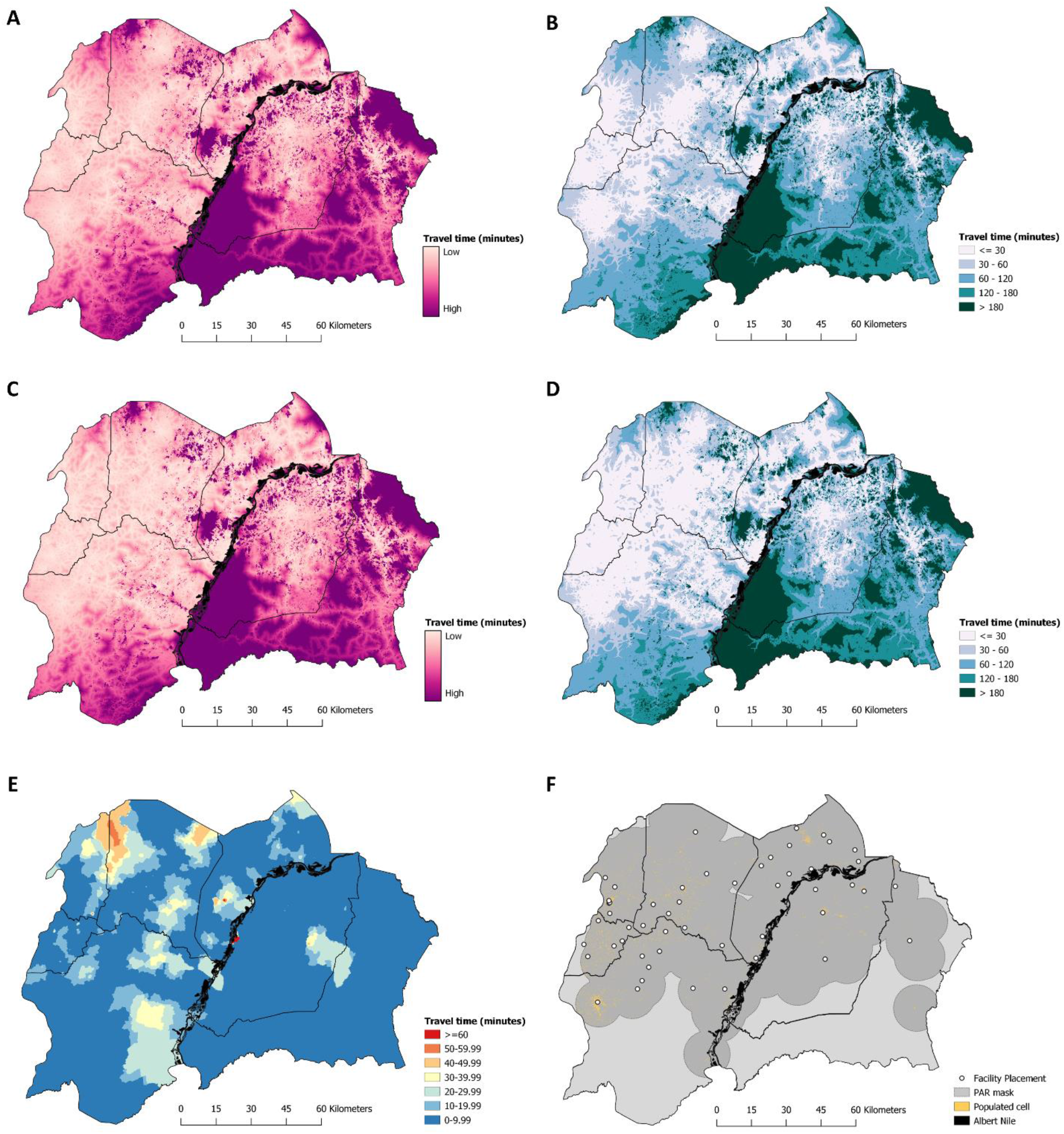
Predicted travel time, in minutes, to a g-HAT diagnostic facility. A) Predicted travel time (in minutes) to reach a diagnostic facility within north-western Uganda, assuming the comprehensive distribution of 170 facilities. B) The predicted travel time surface detailed in A, presented as a categorical surface. C) Predicted travel time (in minutes) to reach a diagnostic facility within north-western Uganda, using the scaled-back number (*n* = 51; (46)) of facilities, with placement derived from the simulation study. D) The predicted travel time surface detailed in C, presented as a categorical surface. E) The difference in travel time, for each gridded cell, between surfaces A and C. Red cells indicate geographic areas where the increase predicted travel time, in the event of scale back, is greatest. F) The location of the optimal placement for the 51 facilities. Yellow pixels (30m × 30m) show the realised distribution of the population at risk, where each pixel represents one or more individuals living inside of the mask for the buffered case data (mask shown as dark grey). The black line running through the centre of our study area represents the Albert Nile river.

To demonstrate the ability of this approach to identify the optimal placement of facilities when the facility number is known, either as derived from the above simulation, or pre-defined, we generated 10,000 estimates of travel time to facilities under varying selection combinations of 51 facilities, the number operating from late 2018 onwards (46). For each simulation, we evaluated the percentage of individuals living within each coverage category (i.e. within 1-hour and 5-hours travel of a facility). The optimal placement of the 51 facilities, from the available distribution of 170 operational facilities resulted in no significant difference in the proportion of the PAR living within 1-hours travel of a facility, when compared to the estimate utilising all 170 facilities (all 170 facilities = 96.15%, 51 facilities = 95.25% coverage). As expected, the coverage of the PAR within 1-hour travel, when quantified using the optimal placement of these 51 facilities far exceeds the in-country coverage target of 50%. Although our 10,000 simulations were only a subset of the total possible combination of 51 facilities from a set of 170, (C(170,51)), we show that multiple combinations results in coverage estimates with negligible differences, particularly in reference to the 5-hour threshold (Fig. 6). Of our 10,000 simulations, 9920 (99.2%) are estimated to have a 5-hour threshold coverage above 99.7% (meeting the coverage value if all 170 facilities are considered). Of these, 9028 (90.28%) have a 1-hour threshold coverage above 90%, whereas the 1-hour coverage is 96.15% if all 170 facilities are considered. We know that the maximum possible coverage cannot exceed that obtained using all 170 facilities (96.15%). As a result, while there may exist an excluded combination that exceeds the coverage obtained by the best identified set in our simulation (95.25%), any improvement would be marginal and negligible (at most 0.9%).

**Figure 6.**
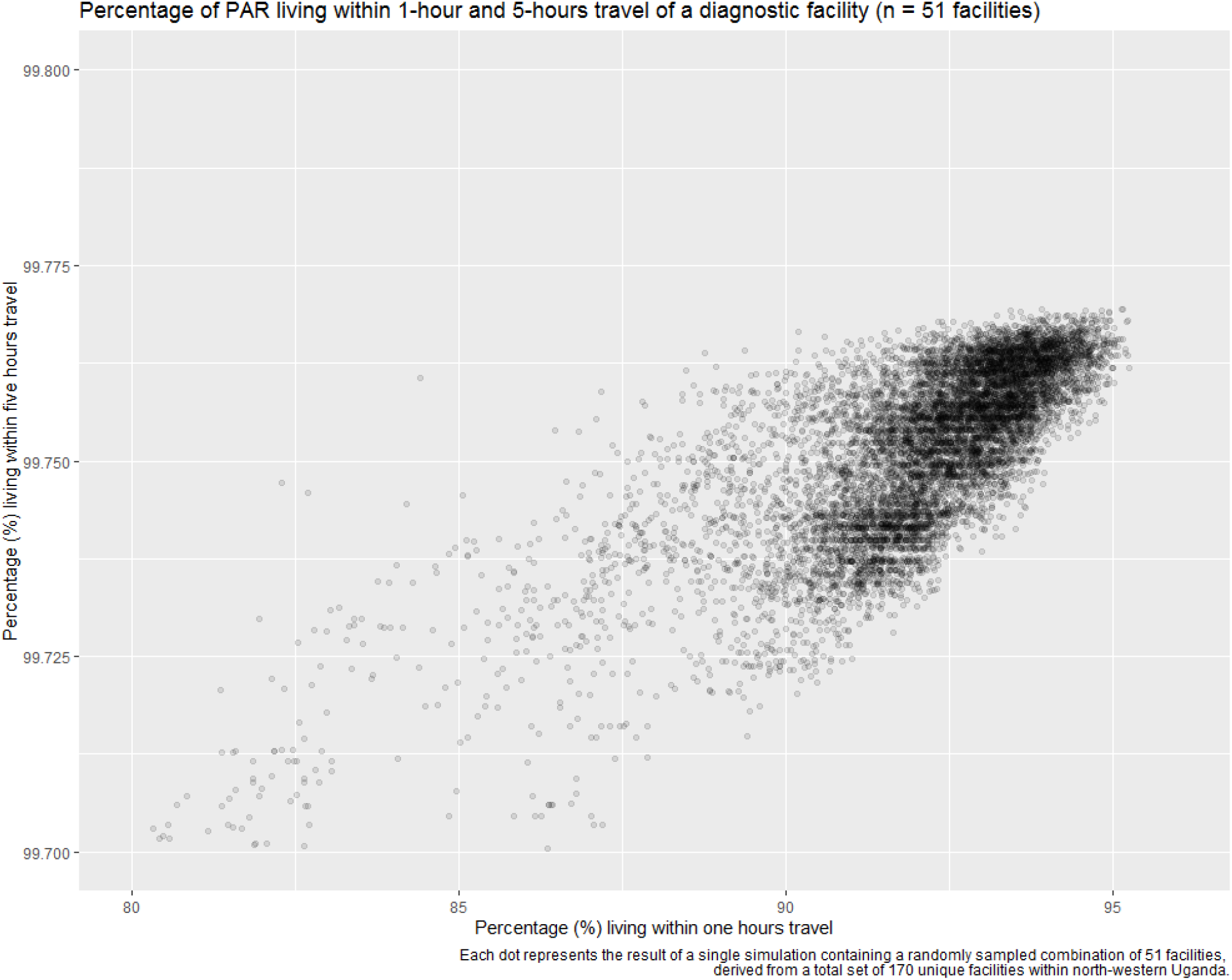
Result of the simulation to determine the optimal combination of 51 facilities, from a total of 10,000 combinations. Each dot represents the result of a single simulation containing a randomly sampled combination of 51 facilities, derived from a total set of 170 unique facilities within north-western Uganda. Due to the majority of simulations far exceeding the 5-hour target, the Y axis is shown as the range 99.700-99.800.

A comparison showing the accessibility to diagnostic facilities is given in Fig. 5, detailing accessibility to all 170 facilities in operation during 2017 (Fig. 5:A-B) and the scaled back number of 51 facilities operating from late 2018 (46), with the optimal placement determined through our approach (Fig. 5:C-D). To aid interpretation, we visualise the cost-distance surfaces on a continuous scale (Fig. 5:A and Fig. 5:C), and on a categorical scale, with travel time binned by 30 minute increments (Fig. 5:B and Fig. 5:D). The difference in travel time from each 30m × 30m gridded cell, between all 170 facilities and 51 facilities is given in Fig. 5:E. A map detailing the optimal placement of the scaled back facilities is provided as Fig. 5:F, and a table listing the name, district and parish of facilities to preserve is provided (Supplementary File 1: Table S2).

Areas of greatest travel time to a facility include those locations which have a low population density, such as Maaji Parish in South-West Adjumani; Waka Parish in Obongi County, Moyo District and Kei Fr Parish in Aringa county, Yumbe District South-West, among others. Additionally, large areas of Southern Arua have populations living ≥1-hour of a diagnostic facility, however, these populations are not generally considered to be living within areas of g-HAT risk so do not affect the population coverage estimates generated here. Maps showing Fig. 5 masked by the population at risk are provided as Supplementary File 1: Figure S2.

## Discussion

Twenty years of a global programme for active screening and treatment of g-HAT has driven the global incidence to a historic low (17). Building on this achievement, recent advances in methods for detecting and treating g-HAT (18, 42), combined with more cost-effective methods of tsetse control (24), are providing the platform for achieving the WHO goals of eliminating g-HAT as a public health problem by 2020 and interrupting transmission completely by 2030. There will be a need for technical guidance on how to monitor the distribution and incidence of g-HAT in this elimination setting. Establishing cost-effective surveillance systems that can be maintained over the next decade will be crucial for the achievement of these ambitious goals.

Using a resistance surface informed by motorbike based travel, and a simulation-based analysis, we determined that the number of operational g-HAT diagnostic facilities within north-western Uganda could be reduced from 170 to three (Fig. 4), whilst ensuring that percentage coverage targets of ≥50% of the PAR living within 1-hours travel and ≥95% 5-hours travel are met. Ensuring that ≥95% of the PAR lives within 5-hours travel of a facility can be achieved by retaining only one well-placed facility. This implies that such targets are easily attainable within the geographic extent of our study area. Scaling back to this degree however poses concerns regarding equity of access (47). For a disease like g-HAT, which predominantly affects rural populations, severe scaling back to achieve moderate targets could leave remote communities without coverage. Ultimately, the degree of scale back should result from discussions with programme implementers and stakeholders, utilising empirical evidence from analyses such as those performed here in addition to the consideration of other operational and contextual factors. Given that it is known that a small number of facilities is enough to achieve the coverage targets, this methodology allows more ambitious targets of coverage to be considered. Specifically, we demonstrate that the number of facilities can be reduced by 70% with minimal loss of coverage.

Through the scale back of the number of facilities, we can improve cost-effectiveness whilst preserving coverage of the entire PAR, including remote populations. These simulations can be easily repeated as the spatial distribution of the PAR changes over time, through substitution of new surfaces detailing contracting or expanding PAR in response to case detection. This will allow a dynamic and rapid response to changing circumstances. The PAR used in this analysis captures the historic distribution of g-HAT within north-western Uganda and is therefore greater than PAR surfaces generated using only contemporary data (e.g. Simarro *et al.* (44)). Our results could however be applied to more recent infection periods.

In this study, national borders, such as those with the neighbouring countries of South Sudan and the Democratic Republic of the Congo, are presumed to prevent individuals who are citizens of either of those two countries from seeking diagnosis within Uganda. Conversely, sub-national borders, (e.g., districts) are presumed not to prevent movement of individuals seeking treatment. As the population density surface used to construct the PAR surface pertains to individuals living within Uganda only, we currently do not capture international movement from neighbouring countries. Our friction surface would need to be expanded to include areas within DRC or South Sudan if we wish to include these populations in our simulations. The methodology described here, although applied to the surveillance of HAT, can also be adapted for a range of elimination scenarios and can also be utilised for the monitoring of emerging diseases as part of national early-warning systems.

Accessibility to a healthcare facility does not correlate precisely with Euclidean distance (48). The use of a validated resistance surface accounts for varying land cover and road network accessibility, factors ultimately affecting travel time, especially for remote and rural communities. Due to data limitations, associated mostly with the clarity of remotely sensed imagery for characterising off-road travel, our resistance surface is representative of travel during the dry season only. Travel to diagnostic facilities will be more difficult during the wet season, particularly along untarmacked roads. By producing our resistance surface at a high spatial resolution, we also provide a more accurate quantification of travel time to a facility than existing surfaces at coarser granularity (49), giving higher precision when quantifying populations within each travel time category. Whilst the time taken to access a facility likely influences an individual’s decision to seek treatment, additional factors are also at play. Future efforts should investigate the use of spatial interaction models to quantify the influence of other metrics on the probability of treatment seeking at a specific location, i.e. gravity models weighted by staff capacity and their levels of training, status of the laboratory and available materials for instance (50, 51). The construction and validation of such models requires empirical measurements of treatment-seeking behaviour (52). Our analysis investigated the optimal retention of facilities in a specific spatial setting. The quality of services available at some facilities will be different to others, however, our approach is sufficiently flexible to allow users to retain required sites (for example, specialist referral centres), and there is scope for national health systems to reallocate resources based on the spatial selection identified.

Although the travel time targets utilised within this analysis were pre-defined (16), further analysis is required to determine the effects of these targets on treatment seeking behaviours. For instance, the cost of continuous travel to a facility which is 1-hour or more from an individual’s residence may present significant barriers to treatment (32), and our analysis shows, that within north-western Uganda, ≥95% of the population are within 5-hours travel to just one well-placed facility, questioning the value of this target for this area. Reducing the number of accessible facilities may also influence factors such as compliance with post-treatment follow up, with a previous study identifying costs for transportation to a facility as a reason for non-completion of treatment (53). Alongside any adopted scale-back approaches, there is a need for sensitisation and information campaigns to inform the PAR of the reduced availability of diagnostics as this information may influence treatment seeking behaviours (54); this campaign can be used alongside a referral system to address such issues.

Scale back of disease surveillance should be integrated within a national development plan and combined with the scale back of intervention programmes. In the context of Uganda, work is ongoing to investigate the effect of reducing the geographical extent of tsetse control operations to areas where g-HAT persists. As disease prevalence changes within a geographic area, approaches are required to ensure that appropriate surveillance and treatment systems are in place. The method described here aims to ensure maximal coverage of surveillance under the most cost-effective scenarios. For national programmes to adopt an optimised approach and integrate this approach within health care delivery systems, it is important to demonstrate that the PAR remains adequately covered, health facilities are accessible, and that a functional referral system is maintained. Providing strategic health facilities with a full range of tests for disease confirmation is critical for elimination.

For other NTDs, plans to scale back surveillance require careful consideration of the country and disease in question, alongside additional context-specific factors (1). As such, the altering of disease surveillance systems should be an iterative process requiring regular reassessment of objectives and methods (9). The approach described here provides a robust, reproducible framework to ensure maximal coverage of the PAR through the minimal number of sites within a passive disease surveillance system.

## Acknowledgements

The data presented in this analysis are based on HAT control activities implemented by the Ugandan Ministry of Health, and supported by the Foundation for Innovative New Diagnostics (FIND) with support from the Bill & Melinda Gates Foundation, the Republic and Canton of Geneva, the German Federal Ministry of Education and Research through KfW, the Swiss Government, UK aid from the UK government and the Medical Research Council. We are grateful to all health facility staff, district leadership and district coordinators in Uganda that enabled this project. The World Health Organization (WHO) have supported targeted active screening in affected villages and kindly provided case data through the human African trypanosomiasis atlas. We also wish to thank Dr Simon Wagstaff and Mr Andrew Bennett (both LSTM) for providing the computational resources for this analysis.

## Author contributions

All authors conceived and planned the study. JL wrote the computer code and designed and did the analyses with input from PRB and MCS. Data was curated by JL, CW and PRB. SJT and MCS provided intellectual input into aspects of this study. All authors contributed to the interpretation of the results. JL wrote the first draft of the manuscript and all authors contributed to subsequent revisions.

## Supplementary Files

Supplementary File 1: Additional methods descriptions and supporting figures and tables. Provided in .docx file format readable by most word processing software.

Supplementary File 2: Population at risk surfaces used within the analysis (north-western Uganda), provided in GeoTiff format at a 30m × 30m spatial resolution. The file may be viewed within open source GIS software such as QGIS and R. Supplementary File 3: Resistance surface used within the analysis (north-western Uganda), provided in GeoTiff format at a 30m × 30m spatial resolution. The file may be viewed within open source GIS software such as QGIS and R.

## References

1. Groseclose SL, Buckeridge DL. Public Health Surveillance Systems: Recent Advances in Their Use and Evaluation. Annual Review of Public Health. 2017;38(1):57–79.

2. German RR, Horan JM, Lee LM, Milstein B, Pertowski CA. Updated guidelines for evaluating public health surveillance systems; recommendations from the Guidelines Working Group. 2001.

3. Choi BC. The past, present, and future of public health surveillance. Scientifica. 2012;2012:875253.

4. Bessell PR, Lumbala C, Lutumba P, Baloji S, Bieler S, Ndung’u JM. Cost-effectiveness of using a rapid diagnostic test to screen for human African trypanosomiasis in the Democratic Republic of the Congo. PLoS One. 2018;13(9):e0204335.

5. Loevinsohn B, Aylward B, Steinglass R, Ogden E, Goodman T, Melgaard B. Impact of targeted programs on health systems: a case study of the polio eradication initiative. American journal of public health. 2002;92(1):19–23.

6. Henderson DA. Surveillance of Smallpox. International Journal of Epidemiology. 1976;5(1):19–28.

7. Reich NG, Lessler J, Varma JK, Vora NM. Quantifying the Risk and Cost of Active Monitoring for Infectious Diseases. Scientific Reports. 2018;8(1):1093.

8. Sutherland CS, Tediosi F. Is the elimination of ‘sleeping sickness’ affordable? Who will pay the price? Assessing the financial burden for the elimination of human African trypanosomiasis *Trypanosoma brucei gambiense* in sub-Saharan Africa. BMJ Global Health. 2019;4(2):e001173.

9. Teutsch SM, Thacker SB. Planning a public health surveillance system. Epidemiological bulletin. 1995;16(1):1–6.

10. Tyson S, Biellik R. Implications for Global Health. In: Cochi SL, Dowdle WR, editors. Disease Eradication in the 21st Century. Cambridge: Cambridge; 2011.

11. Fullman N, Yearwood J, Abay SM, Abbafati C, Abd-Allah F, Abdela J, et al. Measuring performance on the Healthcare Access and Quality Index for 195 countries and territories and selected subnational locations: a systematic analysis from the Global Burden of Disease Study 2016. The Lancet. 2018;391(10136):2236–71.

12. Dickinson B. London Declaration on Neglected Tropical Diseases. Uniting to Combat NTDs.; 2012. Contract No.: 15th July 2019.

13. Regmi S, Callender T, Knox AF, Bhopal A. Research and development funding for 13 neglected tropical diseases: an observational economic analysis. The Lancet. 2014;384:S20.

14. Horlick G, O’Connor J. The Legal Basis for Public Health Surveillance. Concepts and Methods in Infectious Disease Surveillance 2014. p. 7–13.

15. Jajosky RA, Ward J. National, State, and Local Public Health Surveillance Systems. Concepts and Methods in Infectious Disease Surveillance 2014. p. 14–25.

16. NTD Modelling Consortium Discussion Group on Gambiense Human African Trypanosomiasis. Insights from quantitative and mathematical modelling on the proposed 2030 goal for gambiense human African trypanosomiasis (gHAT) [version 1; peer review: 1 approved, 1 approved with reservations]. Gates Open Research. 2019;3(1553).

17. Franco JR, Cecchi G, Priotto G, Paone M, Diarra A, Grout L, et al. Monitoring the elimination of human African trypanosomiasis: Update to 2014. PLOS Neglected Tropical Diseases. 2017;11(5):e0005585.

18. Kennedy PGE. Clinical features, diagnosis, and treatment of human African trypanosomiasis (sleeping sickness). The Lancet Neurology. 2013;12(2):186–94.

19. Franco JR, Simarro PP, Diarra A, Jannin JG. Epidemiology of human African trypanosomiasis. Clinical epidemiology. 2014;6:257–75.

20. Franco JR, Cecchi G, Priotto G, Paone M, Diarra A, Grout L, et al. Monitoring the elimination of human African trypanosomiasis at continental and country level: Update to 2018. PLOS Neglected Tropical Diseases. 2020;14(5):e0008261.

21. World Health Organisation. Global Health Observatory data repository: Human African Trypanosomiasis 2019 [Available from: http://apps.who.int/gho/data/node.main.A1635?lang=en.

22. Checchi F, Cox AP, Chappuis F, Priotto G, Chandramohan D, Haydon DT. Prevalence and under-detection of gambiense human African trypanosomiasis during mass screening sessions in Uganda and Sudan. Parasites & Vectors. 2012;5(1):157.

23. Selby R, Wamboga C, Erphas O, Mugenyi A, Jamonneau V, Waiswa C, et al. Gambian human African trypanosomiasis in North West Uganda. Are we on course for the 2020 target? PLoS neglected tropical diseases. 2019;13(8):e0007550–e.

24. Tirados I, Esterhuizen J, Kovacic V, Mangwiro TNC, Vale GA, Hastings I, et al. Tsetse Control and Gambian Sleeping Sickness; Implications for Control Strategy. PLOS Neglected Tropical Diseases. 2015;9(8):e0003822.

25. Wamboga C, Matovu E, Bessell PR, Picado A, Biéler S, Ndung’u JM. Enhanced passive screening and diagnosis for gambiense human African trypanosomiasis in north-western Uganda – Moving towards elimination. PLOS One. 2017;12(10):e0186429.

26. Checchi F, Filipe JAN, Haydon DT, Chandramohan D, Chappuis F. Estimates of the duration of the early and late stage of gambiense sleeping sickness. BMC Infectious Diseases. 2008;8(1):16.

27. Mehlitz D, Molyneux DH. The elimination of *Trypanosoma brucei gambiense?* Challenges of reservoir hosts and transmission cycles: Expect the unexpected. Parasite Epidemiology and Control. 2019;6:e00113.

28. Facebook Connectivity Lab, Center for International Earth Science Information Network. High Resolution Settlement Layer (HRSL). Source Imagery © 2016 DigitalGlobe, Inc.; 2018.

29. Douglas DH. Least-cost Path in GIS Using an Accumulated Cost Surface and Slopelines. Cartographica: The International Journal for Geographic Information and Geovisualization. 1994;31(3):37–51.

30. OpenStreetMap contributors. Geofabrik OpenStreetMap Data Extracts: 9th July 2019: OpenStreetMap contributors,; 2019 [

31. Longbottom J, Krause A, Torr SJ, Stanton MC. Quantifying geographic accessibility to improve cost-effectiveness of entomological monitoring. PLOS Neglected Tropical Diseases. 2020;14(3):e0008096.

32. Sacks E, Vail D, Austin-Evelyn K, Greeson D, Atuyambe LM, Macwan’gi M, et al. Factors influencing modes of transport and travel time for obstetric care: a mixed methods study in Zambia and Uganda. Health Policy and Planning. 2015;31(3):293–301.

33. U.S. Geological Survey (USGS). Landsat 8 (L8) Data Users Handbook. Sioux Falls, South Dakota: U.S. Geological Survey (USGS),; 2019.

34. Sellers PJ. Canopy reflectance, photosynthesis and transpiration. International Journal of Remote Sensing. 1985;6(8):1335–72.

35. Myneni RB, Hall FG, Sellers PJ, Marshak AL. The interpretation of spectral vegetation indexes. IEEE Transactions on Geoscience and Remote Sensing. 1995;33(2):481–6.

36. Houben RM, Van Boeckel TP, Mwinuka V, Mzumara P, Branson K, Linard C, et al. Monitoring the impact of decentralised chronic care services on patient travel time in rural Africa--methods and results in Northern Malawi. International journal of health geographics. 2012;11:49.

37. Soule RG, Goldman RF. Terrain coefficients for energy cost prediction. Journal of applied physiology. 1972;32(5):706–8.

38. Google Maps. Google Maps Platform: Distance Matrix API 2019 [Available from: https://developers.google.com/maps/documentation/distance-matrix/start.

39. Bisser S, Lumbala C, Nguertoum E, Kande V, Flevaud L, Vatunga G, et al. Sensitivity and Specificity of a Prototype Rapid Diagnostic Test for the Detection of Trypanosoma brucei gambiense Infection: A Multi-centric Prospective Study. PLOS Neglected Tropical Diseases. 2016;10(4):e0004608.

40. Bieler S, Matovu E, Mitashi P, Ssewannyana E, Bi Shamamba SK, Bessell PR, et al. Improved detection of Trypanosoma brucei by lysis of red blood cells, concentration and LED fluorescence microscopy. Acta tropica. 2012;121(2):135–40.

41. Büscher P, Mumba Ngoyi D, Kaboré J, Lejon V, Robays J, Jamonneau V, et al. Improved Models of Mini Anion Exchange Centrifugation Technique (mAECT) and Modified Single Centrifugation (MSC) for Sleeping Sickness Diagnosis and Staging. PLOS Neglected Tropical Diseases. 2009;3(11):e471.

42. Hayashida K, Kajino K, Hachaambwa L, Namangala B, Sugimoto C. Direct blood dry LAMP: a rapid, stable, and easy diagnostic tool for Human African Trypanosomiasis. PLoS Negl Trop Dis. 2015;9(3):e0003578.

43. Franco JR, Cecchi G, Priotto G, Paone M, Diarra A, Grout L, et al. Monitoring the elimination of human African trypanosomiasis: Update to 2016. PLoS Negl Trop Dis. 2018;12(12):e0006890.

44. Simarro PP, Cecchi G, Franco JR, Paone M, Diarra A, Ruiz-Postigo JA, et al. Estimating and Mapping the Population at Risk of Sleeping Sickness. PLOS Neglected Tropical Diseases. 2012;6(10):e1859.

45. Chipeta M, Terlouw D, Phiri K, Diggle P. Inhibitory geostatistical designs for spatial prediction taking account of uncertain covariance structure. Environmetrics. 2017;28(1):e2425.

46. Ndung’u JM, Boulangé A, Picado A, Mugenyi A, Mortensen A, Hope A, et al. Trypa-NO! contributes to the elimination of gambiense human African trypanosomiasis by combining tsetse control with “screen, diagnose and treat” using innovative tools and strategies. PLOS Neglected Tropical Diseases. 2020;14(11):e0008738.

47. Iyer HS, Flanigan J, Wolf NG, Schroeder LF, Horton S, Castro MC, et al. Geospatial evaluation of trade-offs between equity in physical access to healthcare and health systems efficiency. BMJ Global Health. 2020;5(10):e003493.

48. Etherington TR. Least-Cost Modelling and Landscape Ecology: Concepts, Applications, and Opportunities. Current Landscape Ecology Reports. 2016;1(1):40–53.

49. Simarro PP, Cecchi G, Franco JR, Paone M, Diarra A, Ruiz-Postigo JA, et al. Mapping the capacities of fixed health facilities to cover people at risk of gambiense human African trypanosomiasis. International journal of health geographics. 2014;13(1):4.

50. Alegana VA, Wright JA, Pentrina U, Noor AM, Snow RW, Atkinson PM. Spatial modelling of healthcare utilisation for treatment of fever in Namibia. International journal of health geographics. 2012;11:6–.

51. Bakeera SK, Wamala SP, Galea S, State A, Peterson S, Pariyo GW. Community perceptions and factors influencing utilization of health services in Uganda. International Journal for Equity in Health. 2009;8(1):25.

52. Fotheringham AS. Spatial Interaction Models. In: Smelser NJ, Baltes PB, editors. International Encyclopedia of the Social & Behavioral Sciences. Oxford: Pergamon; 2001. p. 14794–800.

53. Lee SJ, Palmer JJ. Integrating innovations: a qualitative analysis of referral non-completion among rapid diagnostic test-positive patients in Uganda’s human African trypanosomiasis elimination programme. Infect Dis Poverty. 2018;7(1):84–-.

54. Wakefield MA, Loken B, Hornik RC. Use of mass media campaigns to change health behaviour. Lancet. 2010;376(9748):1261–71.

