## Supplementary File 1 for "Optimising passive surveillance of a neglected tropical disease in the era of elimination: A modelling study"

### **Methods supplement**

#### **High-resolution accessibility surface**

To quantify travel time to diagnostic facilities, we first sought to generate a friction surface for north-western Uganda. Friction surfaces contain estimates of associated travel cost for gridded cells within a Cartesian plane, and are used within cost-distance analyses, which identify the cumulative cost of traversing each cell based on the given resistance surface and origin (or destination) locations. To construct the resistance surface, data from a variety of sources were pooled, as per the steps below.

##### **Road network**

Shapefiles (geospatial vector data) detailing mapped roads hosted by OpenStreetMap were retrieved from Geofabrik OSM Data Extracts on July 15^th^, 2019. Roads within this source are categorised based off of their social and economic importance, and as such, are classified as being either primary, secondary, tertiary, unclassified, residential, service, track or path when digitised (1). A previous study compared the completeness of OSM data with roads digitized from 3m resolution remotely sensed imagery, identifying a 97.43% agreement between sources within Koboko district, Uganda (2). The referenced study provided confidence in the completeness of the network across the region.

##### **Off-road resistance values**

High-resolution (30m × 30m) remotely sensed imagery for the West Nile Region were obtained from the Landsat 8 satellite (3). The imagery consisted of four tiles for the West Nile Region captured between the 25^th^ January and 1^st^ February 2018, containing minimal cloud cover. Separate tiles were merged to create one scene, utilising ArcGIS version 10.4 (4). The mosaicked image was clipped to administrative boundaries for north-western Uganda. A normalised difference vegetation index (NDVI) was calculated using the processed imagery (5, 6), and off-road resistance values were assigned for different vegetation densities as per a previously described methodology (2). Resistance values represent the time taken (in seconds) to cross each 30m × 30m gridded cell.

##### **Tracks**

To obtain data informing variation in speeds along road class, technicians making routine visits to tsetse traps within Arua, Maracha, Koboko and Yumbe districts, north-western Uganda, were provided with GPS devices. The recording of GPS tracks was performed during February-April 2018 and were representative of travel during the dry season. Trap attendants operate using motorbikes, therefore, observed speeds were characteristic of motorbike-based travel. Devices were configured to record track points at ~15-second intervals. Further detail can be found in Longbottom *et al*. 2020 (2).

##### **Speeds**

Tracking points were converted to polylines, consisting of line segments constructed from five trailing points. These segments were assigned a mean observed speed by calculating the Euclidean distance of each segment and incorporating start and end times. Tracks were overlaid on top of the OSM data, and were classified in the same manner, with segments assigned as being primary, secondary, tertiary or unclassified. These segments were then used to derive a mean observed speed for each OSM road class (Table S1).

##### **Table S1: Speeds assigned to different OpenStreetMap classified roads.**

| OpenStreetMap classification | Mean speed (km/hr) observed | Number of segments |
| --- | --- | --- |
| Primary | 55.95 | 325 |
| Secondary | 53.50 | 1149 |
| Tertiary | 50.57 | 1468 |
| Unclassified | 37.08 | 815 |

##### **Resistance surface**

The updated road network, featuring a cell crossing time based on assigned speeds (representative of on-road resistance), was combined with the NDVI off-road surface to produce one surface detailing associated travel cost (travel time in seconds) for each 30m × 30m cell within north-western Uganda.

##### **Validation**

To assess the accuracy of the resistance surface, a random sample of 1000-paired locations were generated across north-western Uganda. Each pair consisted of an ‘origin’ and ‘destination’ coordinate in decimal degrees, with each ‘origin’ and ‘destination’ being drawn from one of 500 randomised ~2.5km^2^ gridded cells across the study extent. In these selection of these validation points, we assume that people will not cross river Nile in seeking diagnosis, so origin and destination sites occur on the same side of the river. Utilising the ‘gDistance R’ package (7), and the resistance surface, the travel time from each origin and destination was computed. This resulted in a modelled estimate of travel along the resistance surface. To compare performance, the predicted travel time between the same origin and destination locations was obtained from Google Maps (8), using the distance matrix API, facilitated through use of the ‘mapsapi’ R package (9).

#### **Simulation to determine inhibitory sampling distance**

When generating random samples of facilities for use within the simulation to identify the minimum number of required facilities, we adopted a discrete inhibitory sampling approach, as described by Chipeta et al. (10). To determine the minimum distance to utilise within this inhibitory sampling approach, we performed a simulation where we produced a random subsample of facilities from the set available and calculated the mean distance between facilities in the draw. The value used to inform the inhibitory sample is the value in which the mean distance stabilises when the value of n (number of facilities) increase, as per the following:

Suppose $S_{n}$ is the set of all possible combinations of $n$ facilities from the 170 currently available. We wish to obtain an estimate of the distance between facilities for all possible values of $n$, however once $n>1$, the total number of combinations of $n$ facilities is very large and computationally prohibitive. As such, we adopted a simulation approach, such that for $n=2,..,170$ we generated $\hat{S}_{n}$ which consisted of 1000 random samples of $n$ facilities. For each sample,$i=1,\ldots,1000$ we calculated the mean distance ($D_{n}^{i}$) between the $n$ facilities using the Euclidean distance. From this we can obtain $H_{n}$, the average distance between $n$ facilities, across all 1000 random samples. The inhibitory distance value was derived when there was negligible change in $H_{n}$ across all values of $n$, as defined by $H_{n}-H_{n-1}<0.01$. The value derived from the above simulation was 4068.37m, shown as Figure S1.

##### **Figure S1:** Result of the inhibitory sampling simulation.

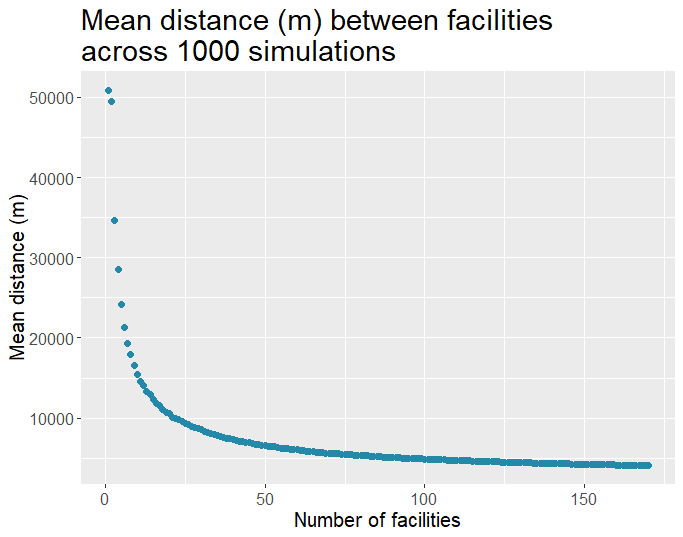

**Table S2: Name and location of spatially optimal facilities to retain.** Facilities identified include private (non-government and faith-based) health centres, and governmental health centres. HCII refers to health centre level II (lowest quality), HCIII refers to health centre level III, HCIV refers to health centre level IV, RMF refers to centres operated by the Real Medicine Foundation and IRC refers to centres operated by the International Rescue Committe.

| Facility name | District | County | Sub-County | Parish | Latitude | Longitude |
| --- | --- | --- | --- | --- | --- | --- |
| Afoji HCII | Moyo | West Moyo | Moyo | Logoba | 3.698040 | 31.683850 |
| Ajikoro HCII | Maracha | Maracha | Oleba | Paranga | 3.339260 | 30.914910 |
| Alere HCII | Adjumani | East Moyo | Pacara | Jikwa | 3.460130 | 31.757700 |
| Aliba HCIII | Moyo | Obongi | Aliba | Indilinga | 3.193720 | 31.526490 |
| Ambelecu HCII | Yumbe | Aringa | Odravu | Lui | 3.366730 | 31.187530 |
| Andelizo HCII | Arua | Terego | Oriama | Maraju | 3.158390 | 31.110360 |
| Ariwa HCIII | Yumbe | Aringa | Ariwa | Rigbonga | 3.242740 | 31.395220 |
| Arra-Pakarukwe HCII | Moyo | West Moyo | Dufile | Arra | 3.613900 | 31.910480 |
| Arua Regional Hospital | Arua | Arua Municipality | Municipality | Municipality | 3.022860 | 30.912040 |
| Atiak HCIV | Amuru | Kilak | Atiak | Kal | 3.261464 | 32.122844 |
| Belameling HCII | Moyo | Obongi | Itula | Legu | 3.476120 | 31.611310 |
| Bibia HCIII | Amuru | Kilak | Atiak | Bibia | 3.472460 | 32.067370 |
| Bileafe HCIII | Arua | Terego | Uriama | Otumbari | 3.106220 | 31.084720 |
| Carlos Medical Centre | Koboko | Koboko | Koboko TC | Mengo Ward | 3.406917 | 30.956467 |
| Discovery Medical Centre | Adjumani | East Moyo | Adjumani TC | Central | 3.370080 | 31.785900 |
| Dramba HCII | Yumbe | Aringa | Drajini | Aupi | 3.320500 | 31.088640 |
| Dranya HCIII | Koboko | Koboko | Dranya | Aunga | 3.367400 | 30.959310 |
| Dricile HCIII | Koboko | Koboko | Midia | Dricile | 3.470180 | 30.978940 |
| Dufile HCIII | Moyo | West Moyo | Dufile | Lebubu | 3.568930 | 31.924350 |
| Elema HCII | Adjumani | East Moyo | Dzaipi | Miniki | 3.477630 | 31.900470 |
| Eremi HCII | Moyo | West Moyo | Metu | Eremi | 3.645760 | 31.811750 |
| Eria HCIII | Moyo | West Moyo | Moyo | Eria | 3.631280 | 31.656600 |
| Fr Bilibao HCIII | Moyo | West Moyo | Metu | Pameri | 3.671840 | 31.789270 |
| Ibakwe HCII | Moyo | West Moyo | Itula | Palorinya | 3.521100 | 31.655110 |
| Iboa HCII | Moyo | Obongi | Itula | Ubbi | 3.539860 | 31.741230 |
| Igamara HC III RMF | Yumbe | Aringa South | Odravu | Abara | 3.311611 | 31.250111 |
| Kamaka HCIII | Maracha | Maracha | Oluffe | Kamaka | 3.247060 | 30.859010 |
| Kerwa HCII | Yumbe | Aringa | Midigo | Kerwa | 3.683390 | 31.291810 |
| Koboko HCIV | Koboko | Koboko | Koboko TC | Koboko TC | 3.409420 | 30.959370 |
| Koro HC III IRC | Yumbe | Aringa North | Kochi | Ombachi | 3.521389 | 31.334900 |
| Kuluba HCII | Koboko | Koboko | Kuluba | Kuluba | 3.505940 | 30.941960 |
| Lefori HCIII | Moyo | West Moyo | Lefori | Ebwea | 3.584680 | 31.583610 |
| Locomgbo HCII | Yumbe | Aringa | Romogi | Locomgbo | 3.484760 | 31.444210 |
| Lodonga HCIII | Yumbe | Aringa | Drajini | Yiba | 3.402090 | 31.127830 |
| Lopke HCII | Yumbe | Aringa | Lomogi | Lopke | 3.552170 | 31.547290 |
| Maduga HCII | Moyo | Obongi | Gimara | Gopele | 3.277880 | 31.551550 |
| Moli HC II | Yumbe | Aringa South | Odravu | Moli | 3.411806 | 31.229056 |
| Morobo Clinic | Koboko | Koboko | Koboko TC | Teremunga Ward | 3.416967 | 30.956033 |
| Mungula HCIV | Adjumani | East Moyo | Itirikwa | Mungula | 3.189920 | 31.794830 |
| Ndapi HC II | Arua | Terego East | Omugo | Ndapi | 3.220194 | 31.152250 |
| Nyumanzi HCII | Adjumani | East Moyo | Dzaipi | Adjugopi | 3.453700 | 31.945980 |
| Obofia HCII | Arua | Terego | Aiivu | Otrevu | 3.201490 | 31.085180 |
| Ocea HCII | Arua | Madi-Okollo | Rigbo | Katiku | 3.076610 | 31.282210 |
| Odupi HCIII | Arua | Terego | Odupi | Ombokoro | 3.296920 | 31.175910 |
| Olujobo HCIII | Arua | Madi-Okollo | Rigbo | Aliba | 3.073740 | 31.405560 |
| Oluvu HCIII | Maracha | Maracha | Oluvu | Ombachi | 3.199980 | 30.874090 |
| Pioneer Medical Centre | Yumbe | Aringa | Yumbe TC | Yumbe TC | 3.467283 | 31.241150 |
| St. Francis Ocodri HCIII | Arua | Terego | Bileafe | Ajiraku | 3.074100 | 31.066870 |
| Tara HCIII | Maracha | Maracha | Tara | Pajama | 3.312780 | 31.033590 |
| Wadra HCIII | Maracha | Maracha | Yivu | Okuvu | 3.260210 | 31.007800 |
| Yivu Abea HCII | Maracha | Maracha | Yivu | Alarapi | 3.240960 | 30.975820 |
| Facility name | District | County | Sub-County | Parish | Latitude | Longitude |
| Afoji HCII | Moyo | West Moyo | Moyo | Logoba | 3.698040 | 31.683850 |
| Ajikoro HCII | Maracha | Maracha | Oleba | Paranga | 3.339260 | 30.914910 |

### **Figure S2: Predicted travel time, in minutes, to a g-HAT diagnostic facility, masked by the population at risk buffered surface.**

A) Predicted travel time (in minutes) to reach a diagnostic facility within north-western Uganda, assuming the comprehensive distribution of 170 facilities. B) The predicted travel time surface detailed in A, presented as a categorical surface. C) Predicted travel time (in minutes) to reach a diagnostic facility within north-western Uganda, using the scaled-back number (n = 51) of facilities, with placement derived from the simulation study. D) The predicted travel time surface detailed in C, presented as a categorical surface. E) The difference in travel time, for each gridded cell, between surfaces A and C. Red cells indicate geographic areas where the increase predicted travel time, in the event of scale back, is greatest. F) The location of the optimal placement for the 51 facilities.

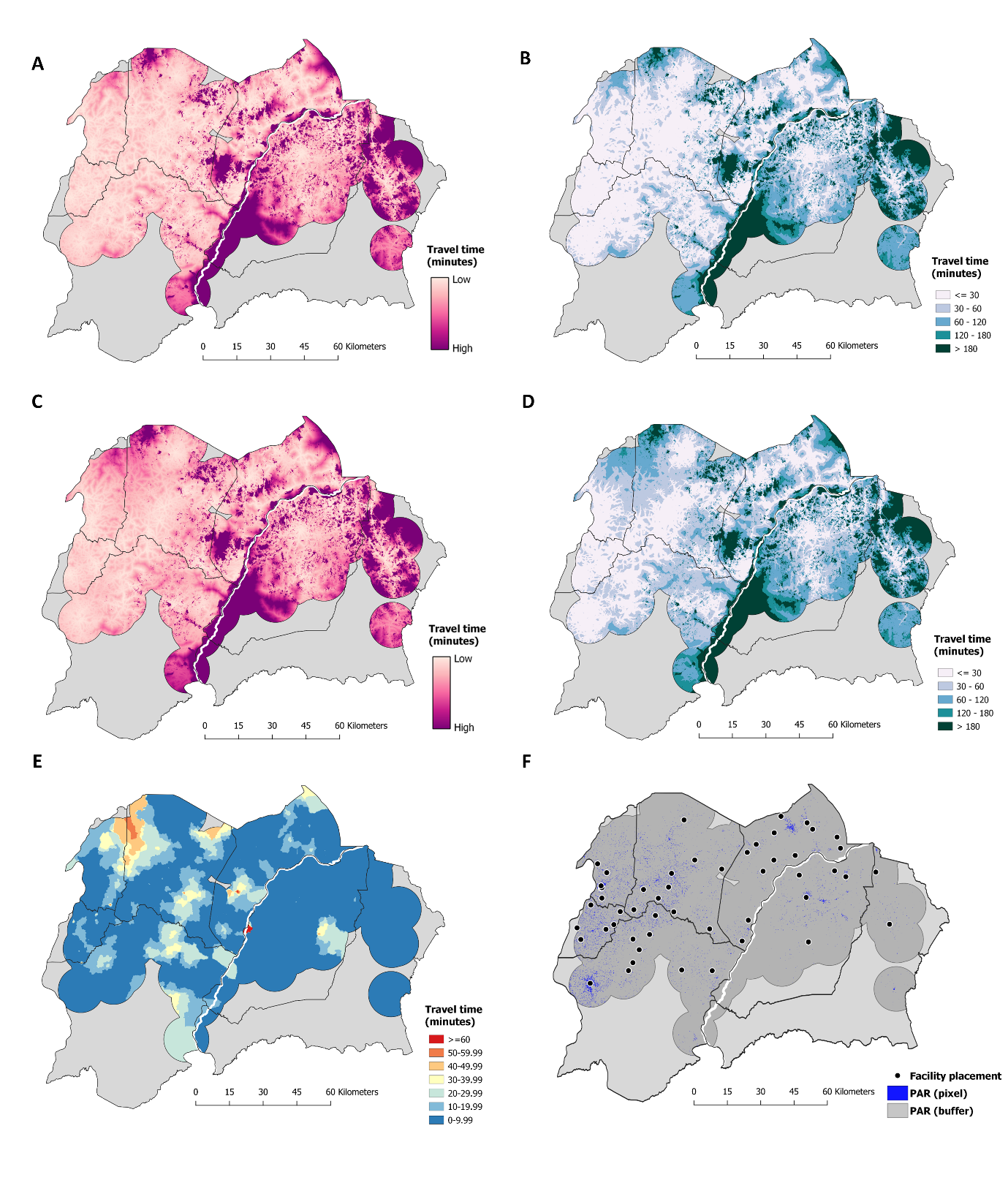

#### **References**

1.	OpenStreetMap. East Africa Tagging Guidelines 2019 [Available from: <https://wiki.openstreetmap.org/wiki/East_Africa_Tagging_Guidelines>.

2.	Longbottom J, Krause A, Torr SJ, Stanton MC. Quantifying geographic accessibility to improve cost-effectiveness of entomological monitoring. PLOS Neglected Tropical Diseases. 2020;14(3):e0008096.

3.	Landsat-8 image courtesy of the U.S. Geological Survey.

4.	ESRI. ArcGIS Desktop: Release 10.4. Redlands, CA; 2018.

5.	Sellers PJ. Canopy reflectance, photosynthesis and transpiration. International Journal of Remote Sensing. 1985;6(8):1335-72.

6.	Myneni RB, Hall FG, Sellers PJ, Marshak AL. The interpretation of spectral vegetation indexes. IEEE Transactions on Geoscience and Remote Sensing. 1995;33(2):481-6.

7.	van Etten J. R Package gdistance: Distances and Routes on Geographical Grids. 2017. 2017;76(13):21.

8.	Google Maps. Google Maps Platform: Distance Matrix API 2019 [Available from: <https://developers.google.com/maps/documentation/distance-matrix/start>.

9.	Michael Dorman, Tom Buckley, Alex Dannenberg, Bhaskar M. mapsapi: 'sf'-Compatible Interface to 'Google Maps' APIs. 2020.

10.	Chipeta M, Terlouw D, Phiri K, Diggle P. Inhibitory geostatistical designs for spatial prediction taking account of uncertain covariance structure. Environmetrics. 2017;28(1):e2425.
